# Transmission network reconstruction for foot-and-mouth disease outbreaks incorporating farm-level covariates

**DOI:** 10.1101/835421

**Authors:** Simon M. Firestone, Yoko Hayama, Max S. Y. Lau, Takehisa Yamamoto, Tatsuya Nishi, Richard A. Bradhurst, Haydar Demirhan, Mark A. Stevenson, Toshiyuki Tsutsui

**Affiliations:** Melbourne Veterinary School, Faculty of Veterinary and Agricultural Sciences, The University of Melbourne, Parkville, VIC 3010, Australia; Viral Disease and Epidemiology Research Division, National Institute of Animal Health, National Agriculture Research Organization, Tsukuba, Ibaraki 305-0856, Japan; Department of Biostatistics and Bioinformatics, Rollins School of Public Health, Emory University, Atlanta, Georgia, United States of America; Exotic Disease Research Station, National Institute of Animal Health, National Agriculture and Food Research Organization, Kodaira, Tokyo, 187-0022, Japan; Centre of Excellence for Biosecurity Risk Assessment, The University of Melbourne, Parkville, VIC 3010, Australia; Mathematical Sciences Discipline, School of Science, RMIT University, Melbourne, VIC 3000, Australia

## Abstract

Transmission network modelling to infer ‘who infected whom’ in infectious disease outbreaks is a highly active area of research. Outbreaks of foot-and-mouth disease have been a key focus of transmission network models that integrate genomic and epidemiological data. The aim of this study was to extend Lau’s systematic Bayesian inference framework to incorporate additional parameters representing predominant species and numbers of animals held on a farm.

Lau’s Bayesian Markov chain Monte Carlo algorithm was reformulated, verified and pseudo-validated on simulated outbreaks populated with demographic data Japan and Australia. The modified model was then implemented on genomic and epidemiological data from the 2010 outbreak of foot-and-mouth disease in Japan, and outputs compared to those from the SCOTTI model implemented in BEAST2.

The modified model achieved improvements in overall accuracy when tested on the simulated outbreaks. When implemented on the actual outbreak data from Japan, infected farms that held predominantly pigs were estimated to have five times the transmissibility of infected cattle farms and be 49% less susceptible. The farm-level incubation period was 1 day shorter than the latent period, the timing of the seeding of the outbreak in Japan was inferred, as were key linkages between clusters and features of farms involved in widespread dissemination of this outbreak. To improve accessibility the modified model has been implemented as the R package ‘BORIS’ for use in future outbreaks.

## Introduction

Outbreaks of foot-and-mouth disease (FMD) in previously free countries cause severe and widespread socio-economic impacts [1]. FMD-free countries therefore have stringent biosecurity measures in place to prevent incursions and investigate outbreaks very thoroughly. Following a review of outbreaks in non-endemic regions covering the period 1992 to 2003 [2], there have been a series of costly outbreaks in previously free countries, including those in the United Kingdom in 2007 [3], Taiwan in 2009 [4], Japan in 2010 [5] and three independent introductions into South Korea between 2010 and 2011 [6]. Many of these outbreaks are detailed in a recent review [7].

The inference of ‘who infected whom’ in infectious disease outbreaks has gained considerable momentum in the wake of rapid advances in genome sequencing [8]. Accurate inference of the transmission network and epidemiological parameters can aide in decision-making in the early phases of an outbreak in numerous ways, including: assisting in targeting who to investigate; uncovering whether unsampled (and possibly as yet undetected) sources are seeding new clusters; and establishing whether or not control measures, as implemented, are effectively breaking transmission. Retrospective reconstruction of outbreak networks is useful for establishing risk factors for transmission and failures in biosecurity, targeting surveillance and planning for how to respond most appropriately to future outbreaks. Bayesian models that combine genomic and epidemiological data to infer the transmission network of outbreaks have been developed for a range of emerging infectious diseases and transboundary animal diseases including highly pathogenic avian influenza [9, 10], Ebola [11] and FMD [10, 12–14]. These have recently been reviewed and benchmarked for application in FMD outbreaks [15]. The best-performing approaches in that previous analyses were Lau’s joint Bayesian inference framework [12], the Structured Coalescent Transmission Tree Inference (SCOTTI) model version 1.1.1 [14] and a modification to Cottam’s original frequentist approach [15, 16]. None of these models include farm-level covariates other than the spatial relationship between farm locations.

In April 2010, an outbreak of FMD was detected in the Miyazaki Prefecture of Japan. This was the first outbreak in the country for 10 years and prior to this outbreak vaccination had not been practiced for FMD in Japan. The earliest detected infected premises (IPs) included mostly beef cattle farms, with rapid spread to pig and dairy cattle farms across the extent of the Prefecture. The outbreak was officially detected on 20 April 2010 based on PCR positive test results on samples from cattle at a fattening farm, though non-specific clinical signs had first been detected, but not diagnosed as FMD, in a cow on this farm on 9 April 2010, and even earlier, on 31 March 2010 in water buffalo on a nearby farm [17]. The outbreak lasted 2.5 months, during which time 292 IPs were detected and around 200,000 infected animals (cattle, pigs, water buffalos, goats and sheep) were culled to contain spread. A further 87,000 animals that were vaccinated during the control program were also slaughtered to expedite the resumption of international trade in livestock produce. Detailed epidemiological descriptions of the outbreak, genomic analyses, risk factor investigations and simulation studies have been published [5, 17–23].

The aim of the present study was to extend Lau’s systematic Bayesian inference framework to incorporate farm-level covariates representing the predominant species and numbers of animals held on infected farms. Specific further objectives included evaluating the performance of the modified model in characterising the transmission process, and estimating key epidemiological and phylogenetic parameters on data from the 2010 FMD outbreak in Japan, alongside other available approaches.

## Materials and Methods

### Model formulation and modification

The model developed here is an adaptation of Lau’s joint Bayesian Markov Chain Monte Carlo (MCMC) inference framework [11, 12]. In Lau’s original model, the total probability of individual *j* becoming infected during time period [*t*, *t* + *dt*] was given by:

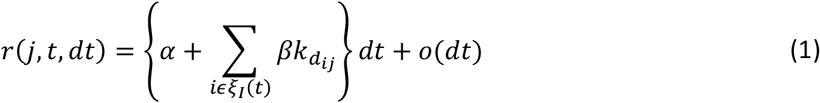

where *ξ*_*I(t)*_ is the set of all infectious premises at time *t*, *α* is the background rate of infection, *β* is the secondary transmission rate, *k*_*dij*_ is a transmission kernel function used to represent the spatial relationship between premises with *o(dt)* representing probability of individual *j* being infected by multiple sources of infection in the small period *dt*, here the power law kernel was assumed of the form:

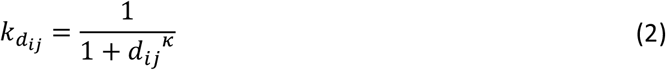

where *d*_*ij*_ is the Euclidean distance between the premises and *κ* is an inferred parameter. Other options for the spatial kernel include exponential, Cauchy and Gaussian decay (not tested here).

In the present analysis, the term *β* in equation (1) was reformulated as *β*_*ij*_ to incorporate additional terms that represent modifications to the transmissibility of each infectious farm, *Inf*_*i*_, and the susceptibility of each susceptible farm, *Susc*_*j*_, such that:

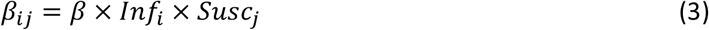

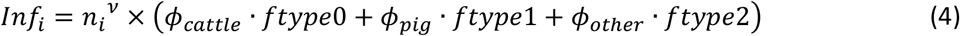

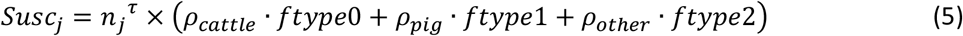

where *n*_*i*_ and *n*_*j*_ represent the number of animals on premises *i* and *j*, respectively, and *ν* and *τ* are inferred parameters that allow for nonlinear effects of holding size [24]. We allowed three levels (modulated by an indicator variable for farm type, *ftype*) for inferred parameters representing the effect of the predominant species on premises *i* and *j* on transmissibility, such that *ϕ*_*pig*_ and *ϕ*_*other*_ represented the component of instantaneous hazard modified by the infectiousness of predominantly pig and other farms (compared to a reference category of predominantly cattle farms, i.e. *ϕ_cattle_=1*), respectively, and *ρ*_*pig*_ and *ρ*_*other*_ represented the susceptibility of predominantly pig and other farms (compared to a reference category of predominantly cattle farms, *ρ_cattle_=1*), respectively. This accounts for a well described biological feature of transmission whereby the minimum infectious doses by inhalation for cattle, sheep and goats are much lower than those of pigs, whereas infectious pigs excrete considerably more virus than these ruminant species [25] and is similar in underlying structure to one of the key simulation models implemented on data from the 2001 FMD outbreak in the United Kingdom [24, 26]. The parameter *β* was retained for scaling purposes. A further modification to the model was also tested, where the infectivity and susceptibility terms were normalised by the population mean infectivity and susceptibility, respectively.

### Model verification and pseudo-validation

The modified model was verified on three FMD outbreak datasets simulated following a previously described approach [27] based on Sellke thresholds [28]. These ‘model verification’ simulation runs (designated J1–J3) were parameterised with the same underlying population structure as areas of Miyazaki Prefecture in Japan from 2010, with differing numbers of susceptible farms and different plausible transmission and genomic parameters.

The modified model was then pseudo-validated by testing on three previously described FMD outbreak datasets simulated in the Australian Animal Disease Simulation (AADIS) model [29], a completely different modelling framework. Corresponding phylogenetic trees nested within the known transmission networks were simulated with VirusTreeSimulator (https://github.com/PangeaHIV/VirusTreeSimulator; last accessed 31 October, 2017) and SeqGen version 1.3.3 [30]. These simulated Australian FMD outbreak datasets were designated A1–A3. All simulated datasets are provided in supplementary materials (S1) along with detailed descriptions of their parameterisation.

### Case study: 2010 outbreak of FMD in Miyazaki Prefecture, Japan

The 2010 Miyazaki FMD outbreak datasets analysed were provided by the National Institute of Animal Health and comprised premises-level covariate data on 292 infected premises and 104 L-fragment consensus nucleotide sequences of virus isolates from animals on these farms, prepared as previously described [5, 18, 20, 21]. Sequences were tested for recombination using RDP4 [31] and for the best fitting DNA substitution model using MEGA version 7.0 [32], as assessed based on the lowest Bayesian Information Criterion.

### Model implementation

The modified joint Bayesian MCMC inference of the transmission tree was implemented on a parallel computing cluster with 4 chains of 1 million iterations, the first 20% of each discarded as burn-in and the remainder thinned by 1000 based on assessment of convergence and autocorrelation, with Gelman and Rubin’s shrink factor [33], visually and by calculation of autocorrelation and effective sample size using Tracer [34]. All unobserved parameters (Table 1) were given uninformative flat priors and imputed as described previously [12]. The MCMC was initialised with a transmission tree with initial sources selected randomly from amongst those estimated to hold infectious animals at the estimated time of exposure of each IP. If there were no potential sources at the estimated time of exposure of an IP the proposed source for this IP was initialised with a value to represent seeding from a non-observed IP. The initiating single universal master sequence was assumed to be the consensus sequence of all available genomic data.

**Table 1:** Key parameters in the Bayesian MCMC inference.

| Parameter | Type | Description |
| --- | --- | --- |
| $t.samp_j, S_j$ | Observed | The timing of sampling and available sequences for infected premises in the dataset. |
| $\psi_j, t_{e_j}, t_{i_j}$ | Latent | The source and timing of exposure and onset of infectiousness for each exposed site $j$. |
| $Gt_j$ | Latent | The sequence on each infected premises at each sampling and transmission time ($t$). |
| $\alpha$ | Latent | The background rate of infection. |
| $\beta, \beta_{ij}$ | Latent | The secondary transmission rate, with and without additional farm-level covariates. |
| $d_{ij}$ | Observed | Euclidean distance between premises $i$ and $j$. |
| $\kappa$ | Latent | The power of the spatial transmission kernel. |
| $n_i, n_j$ | Observed | Number of animals on premises $i$ and $j$. |
| $\nu$ | Latent | The effect (power) of number of animals on premises-level infectivity for farms. |
| $\tau$ | Latent | The effect (power) of number of animals on premises-level susceptibility for farms. |
| $\phi_{cattle}, \phi_{pig}, \phi_{other}$ | Latent | The multiplicative effect of predominant species on premises-level infectivity. |
| $\rho_{pig}, \rho_{other}$ | Latent | The multiplicative effect of predominant species on premises-level susceptibility. |
| $\mu_1, \mu_2$ | Latent | The rates of transitions and transversions. |
| $mean(lat), var(lat)$ | Latent | The mean and variance of the duration of the farm-level latent period. |
| $c$ | Latent | The mean period from onset of infectiousness to the last day of culling (i.e., the farm-level infectious period). |
| $p$ | Latent | Probability that a nucleotide base of a primary sequences differs from that in the universal master sequence. |

### Comparative analyses

The 2010 Miyazaki FMD outbreak dataset was also analysed by preparing temporal transmission windows [16] and inferring the transmission network and phylogenetic parameters with the SCOTTI model version 1.1.1 [14], implemented in BEAST version 2.4.7 [35]. The HKY substitution model [36] was assumed with 2 independent chains of 10 million MCMC iterations, each with 20% discarded as burn-in and thinned by 20000 based on assessment of convergence and autocorrelation. In this coalescent model with migration, each IP was modelled as a ‘host’, each with a distinct diverse pathogen population undergoing genetic evolution. Transmissions between hosts were modelled as ‘migration’ events and the maximum number of hosts was set to 10 times the number of sequences available to allow for unobserved IPs, observed IPs for which genomic data was missing and seeding from external clusters. All unobserved parameters were given uninformative flat priors and the following were inferred: the mutation rate, the ratio of transitions to transversions, the rate of transmission between hosts, the total number of hosts (including non-sampled IPs), the number of pathogen lineages per host and the tree height (from which the delay between origin and detection of the outbreak could be estimated).

The code for implementing the modified Lau model has been incorporated into a freely available R package named Bayesian Outbreak Reconstruction, Inference and Simulation (BORIS) [37]. The descriptive analyses of all model outputs was undertaken in the R statistical package version 3.4.3 [38], using the libraries epiR v0.9-93 [39], statnet v2016.9 [40], coda v0.19-1 [41] and ggplot2 [42]. In all comparisons, model accuracy in inferring the transmission network was considered as the proportion of infected premises for which the true source was the proposed source with the highest posterior probability density [15]. The effect of features of the inferred transmission network on the reproductive number was inferred as previously described [43].

## Results

### Model verification and pseudo-validation

The modified version of the model demonstrated improved performance in each of the simulated model runs (Figure 1 and supplementary materials, S2). Overall accuracy improved by 6.2% in verification runs J1–J3 (range: 5.4–6.9%) and by 4.7% in pseudo-validation runs A1–A3 (range: 2.3– 7.8%). Accuracy improvements occurred over the full range of model support values. Posterior probability density (model support) for proposed sources was higher for outputs from the modified versus the original model for all verification runs (Wilcoxon signed-rank p-values all <0.001) and comparable for pseudo-validation runs; higher support has previously been associated with higher accuracy. The performance of the modified-normalised version was very similar to the modified version without normalisation, with the non-normalised version demonstrating typically 1–2% better accuracy.

**Figure 1:**
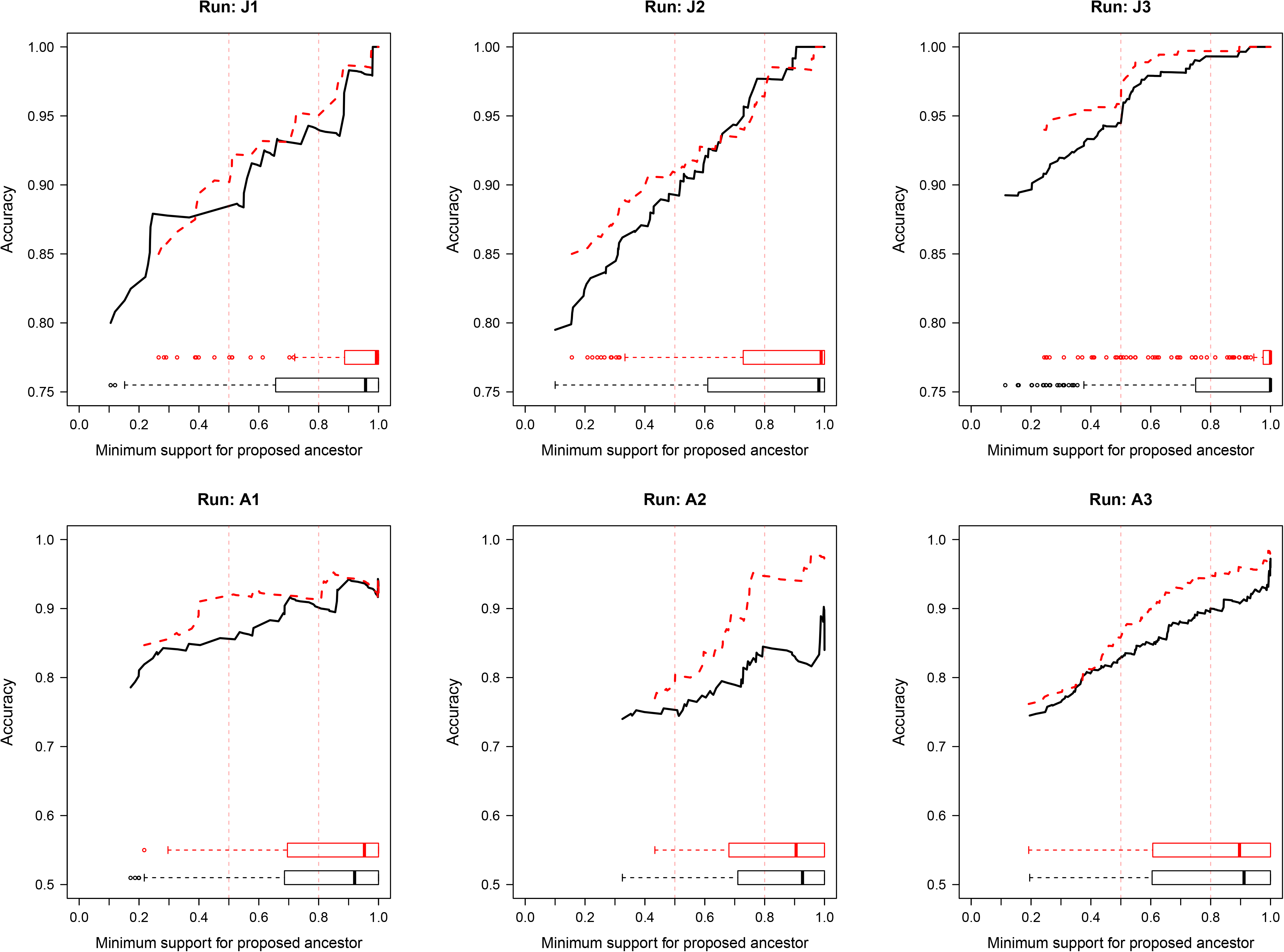
Comparison of the accuracy of inferences of proposed sources of infection for six simulated outbreaks of foot-and-mouth disease in Japan and Australia. Black line = original formulation; red = modified model. Runs J1, J2 and J3 simulated in the same framework as the modified model. Runs A1, A2, A3 simulated in using the Australian Animal Disease Simulation model. Accuracy was defined as the proportion of infected premises for which the true source was the proposed source with the highest posterior probability density. Vertical reference lines denote proposed ancestors with >50% and >80% model support, respectively.

Posterior distributions of the inferred epidemiological and phylogenetic parameters are presented in Supplementary Materials S3 by model run, compared to the known values. In validation runs, the models were highly accurate and comparable in their inferences of *α*, the mutation rate and transition-to-transversion ratio, farm-level latent and infectious periods, the spatial kernel shape parameter (*κ*), and the farm-level transmissibility (*ϕ*) and susceptibility (*ρ*) weighting parameters, and indices for the effects of number of animals per farm (*ν* and *τ*). In runs J1 and J2, the models were highly accurate in their inference of the secondary transmission rate (*β*). In run J3, which had an extremely low value of *β* all of the models overestimated the true value of 6 × 10^−4^. The modified model had the least discrepancy, with its highest probability density region (HPD) ranging from 2 to 3 times the true value, the inferred values for the original and modified-normalised models were out by >200-fold. In pseudo-verification runs, the models were highly accurate and comparable in their inferences of the transition-to-transversion ratio, however all three models underestimated the mutation rate by between 41% and 49%. The rest of the inferred parameters are not directly analogous to those used in the simulation framework for pseudo-validation, so could not be directly compared to known values.

### Case study: 2010 outbreak of FMD in Miyazaki Prefecture, Japan

Each of the 104 sequences were 7667 nucleotides in length, no recombination was detected. The best-fitting nucleotide substitution model was the Tamura-Nei (TN93) model with non-uniformity of the evolutionary rate among sites represented using a discretised Gamma distribution with five categories, an estimated shape parameter of 0.13, assuming that none of the sites were evolutionarily invariable and a transition to transversion ratio of 9.08 (see supplementary materials, S4 for further detailed results).

The transmission network inferred using the modified Lau MCMC algorithm is presented in arbitrary space in Figure 2. Posterior estimates of the key epidemiological and phylogenetic parameters from the modified version of the Lau model are presented in Table 2. Networks for the original and modified-normalised model formulations are provided as Supplementary Materials (S5) highlighting differences to the presented network. The root of the inferred transmission tree was inferred with very high model support. Transmission from an external source was inferred to have most likely occurred 31 days prior to the outbreak being detected (i.e., on 19 March 2010; 95% HPD: 8 and 25 March 2010). At the point of outbreak detection (on 20 April 2010) it was inferred that there were 15 farms already infected. The median diagnostic delay (time from inferred exposure at a farm until day of sampling) was estimated to be 9.7 days (range: 4.6, 32.9 days).

**Table 2:** Epidemiological and phylogenetic parameters inferred for the 2010 outbreak of foot-and-mouth disease in Miyazaki Prefecture, Japan, by transmission network model.

| Model parameter (units) | Lau's joint inference, modified<br>Posterior Median [95% HPD] | *Structured Coalescent Transmission*<br>*Tree Inference* Posterior Median [95%<br>HPD] |
| --- | --- | --- |
| Primary transmission rate, $\alpha$ | $4.0 \times 10^{-5}$ [$0.1 \times 10^{-5}$, $2.1 \times 10^{-4}$] | — |
| Secondary transmission rate, $\beta$ | 0.063 [0.016, 0.142] | — |
| Mutation rate (substitutions site<sup>-1</sup> day<sup>-1</sup>) | $1.83 \times 10^{-5}$ [$1.63 \times 10^{-5}$, $2.06 \times 10^{-5}$] | $2.31 \times 10^{-5}$ [$1.73 \times 10^{-5}$, $2.89 \times 10^{-5}$] |
| Transition to transversion ratio | 6.95 [5.20, 9.57] | 10.12 [6.68, 14.33] |
| Delay from origin of epidemic to outbreak detection (days) | 30.9 [25.9, 42.3] | 38.5 [24.4, 56.5] |
| Effective population size<sup>a</sup> | — | 18.9 [8.6, 34.5] |
| Number of farms infected at outbreak detection | 15 [11, 30] | — |
| Farm-level incubation period (days) | 5.6 [2.6, 13.8] | — |
| Farm-level latent period, $mean(lat)$ (days) | 6.8 [5.2, 8.1] | — |
| Farm-level infectious period, $c$ (days) | 15.2 [13.6, 17.2] | — |
| Spatial kernel scaling parameter, $\kappa$ | 1.79 [1.54, 2.04] | — |
| Infectivity of pig farms vs. cattle farms, $\phi_{pigs}$ | 5.15 [2.64, 11.59] | — |
| Infectivity of other farms vs. cattle farms, $\phi_{other}$ | 0.50 [0.11, 1.67] | — |
| Effect of farm size on infectivity, $\nu$ | 0.08 [0.00, 0.26] | — |
| Susceptibility of pig farms vs. cattle farms, $\rho_{pigs}$ | 0.51 [0.30, 0.83] | — |
| Susceptibility of other farms vs. cattle farms, $\rho_{other}$ | 0.45 [0.14, 1.22] | — |
| Effect of farm size on susceptibility, $\tau$ | 0.23 [0.11, 0.35] | — |

HPD = Highest probability density region; IP = infected premises. <sup>a</sup> Estimated from structured coalescent migratory model based on within-host (here, within-farm) effective population size ( $N_e$ ), migration rate and proportion of hosts with consensus support that their source was sampled.

**Figure 2:**
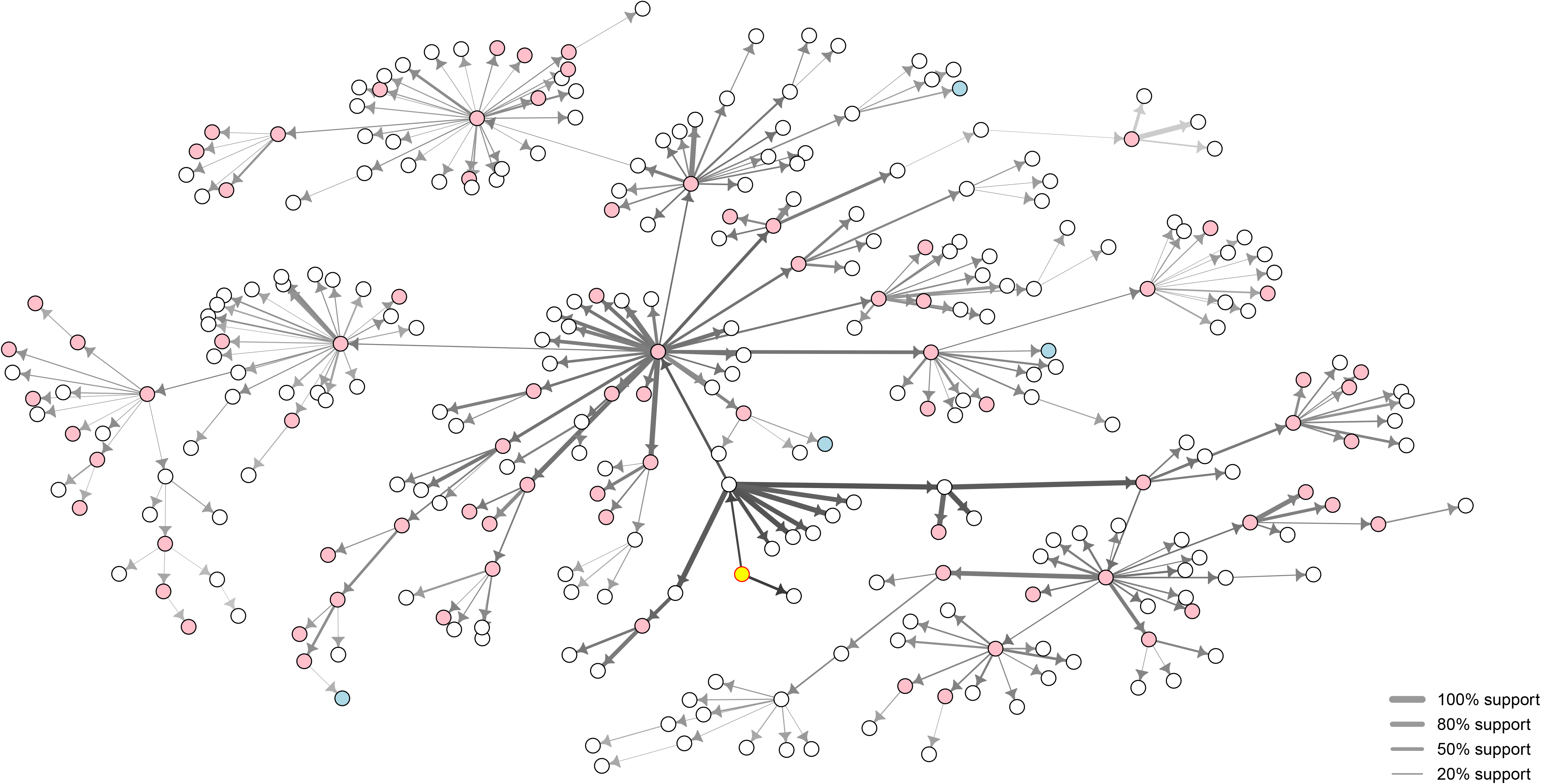
Inferred transmission network for the 2010 outbreak of foot-and-mouth disease in Miyazaki Prefecture, Japan, in arbitrary space. Model support for the proposed ancestor represented by edge width. Darker shading of edges represents earlier inferred transmission events in the outbreak. Farms holding predominantly pigs, cattle and other species are represented by pink, white and blue nodes, respectively. Case numbers randomised for confidentiality.

Of the 292 IPs, only 47 had a proposed source from Lau’s modified algorithm with model support >50%, of these only 18 links had model support >80%. Model support was highest for inferred transmission events earlier in the outbreak (geometric mean support for events in first 4 weeks was 74.8%, whereas for events in the mid and latter 4-week periods of the outbreak geometric mean support was 24.1% and 12.2%, respectively), likely relating to the density of genomic sampling. The longest of the inferred chains of infection involved 8 transmission events, with 93% of transmission chains being ≤5 events in length. The scale-free properties of the transmission network’s out degree distribution (coefficient of variability = 3.3), suggested a multiplying effect on the basic reproductive number of 12.0. The geometric mean number of secondarily infected premises for IPs exposed in the first 4 weeks of the outbreak was 5.9, dropping to 3.2 and 1.3 for IPs exposed in the middle and latter 4-week intervals of the outbreak, respectively. This demonstrates the effectiveness of animal movement controls and other measures.

Farms that kept predominantly pigs were 5.15 times more infectious than cattle farms (Table 2). The eleven farms that were inferred to have led to the highest number of secondary infections were all pig farms. Those farms that predominantly kept other species appeared less infectious than cattle farms, however as there were only five ‘other’ farms the HPD for *ϕ*_*other*_ crossed the null value of 1. Farms that kept predominantly pigs were 49% less susceptible than cattle farms. Those farms that predominantly kept other species were 55% less susceptibility than cattle farms (noting that the HPD again crossed 1, due to low numbers in this group). The number of animals on a farm had more influence on farm-level susceptibility than infectivity.

The posterior estimates of the mean farm-level incubation, latent and infectious periods were 5.9, 6.8 and 15.2 days, respectively. Based on the shape of the inferred spatial transmission kernel (Figure 3), most of the density of risk is within 15 km of an infected premises. Most parameter inferences were highly comparable across model runs (modified versus original and normalised). An exception was the secondary transmission rate (*β*) which from the modified-normalised model outputs was inferred to be an order of magnitude higher than as inferred in the original and modified formulation. The HPDs of most of the inferred parameters overlapped with those used in the model verification runs J1–J3.

**Figure 3:**
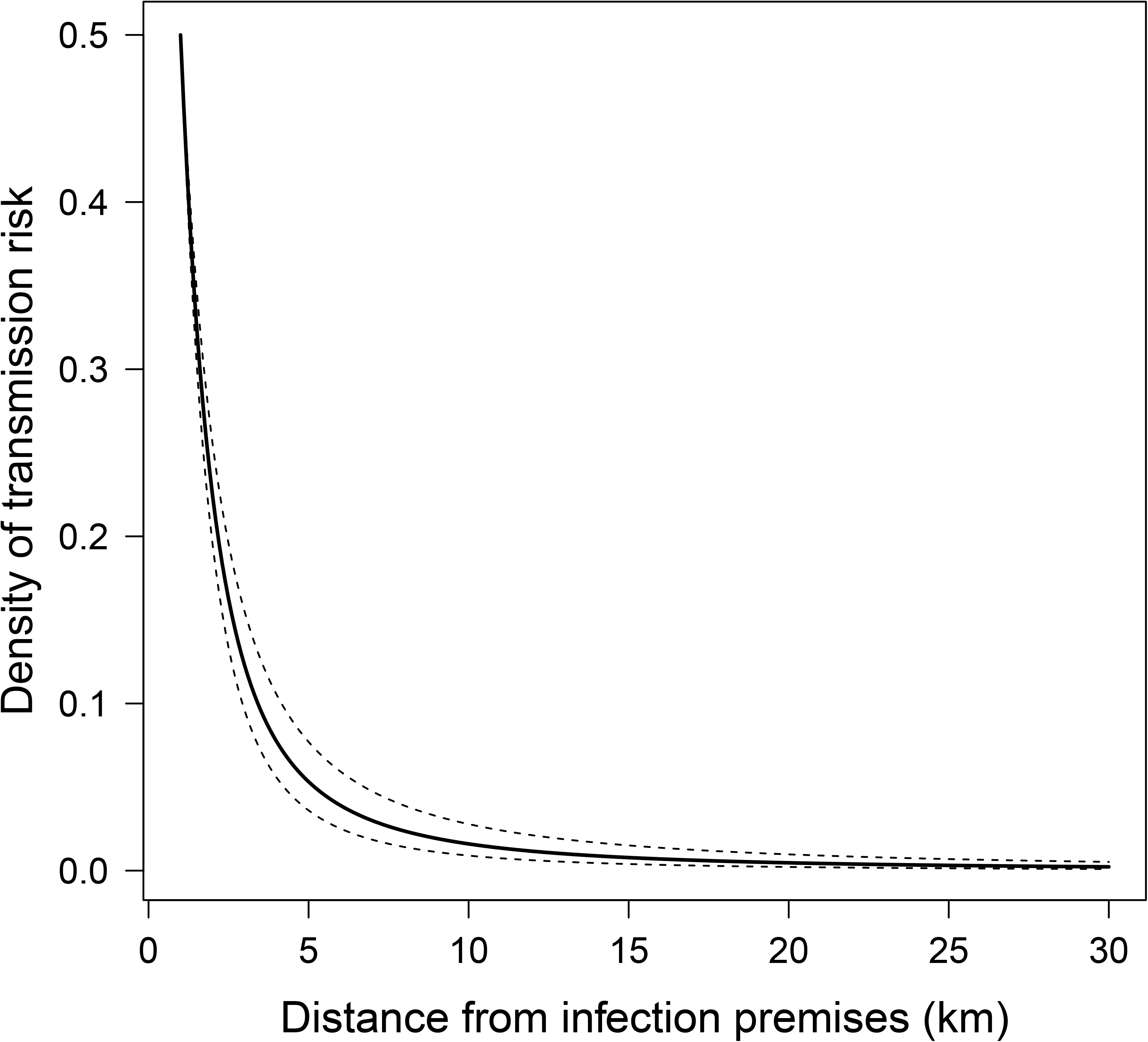
Inferred spatial transmission kernel shape for the 2010 outbreak of foot-and-mouth disease in Miyazaki Prefecture, Japan. Bold line represents posterior median prediction and dashed lines represent 95% highest probability density region.

### Comparative analyses

Transmission windows estimated by Cottam’s approach, are presented for the 20 IPs with earliest dates of onset in Figure 4. Based on this approach, at least ten IPs had already been exposed by the time the outbreak was detected. There were only seven IPs for which the Lau modified and SCOTTI models agreed on source. Amongst the 104 IPs for which genomic data were available, proposed sources for 13 IPs inferred by the SCOTTI algorithm were on the transmission pathways inferred by the Lau model (which included both sampled and unsampled sources).

**Figure 4:**
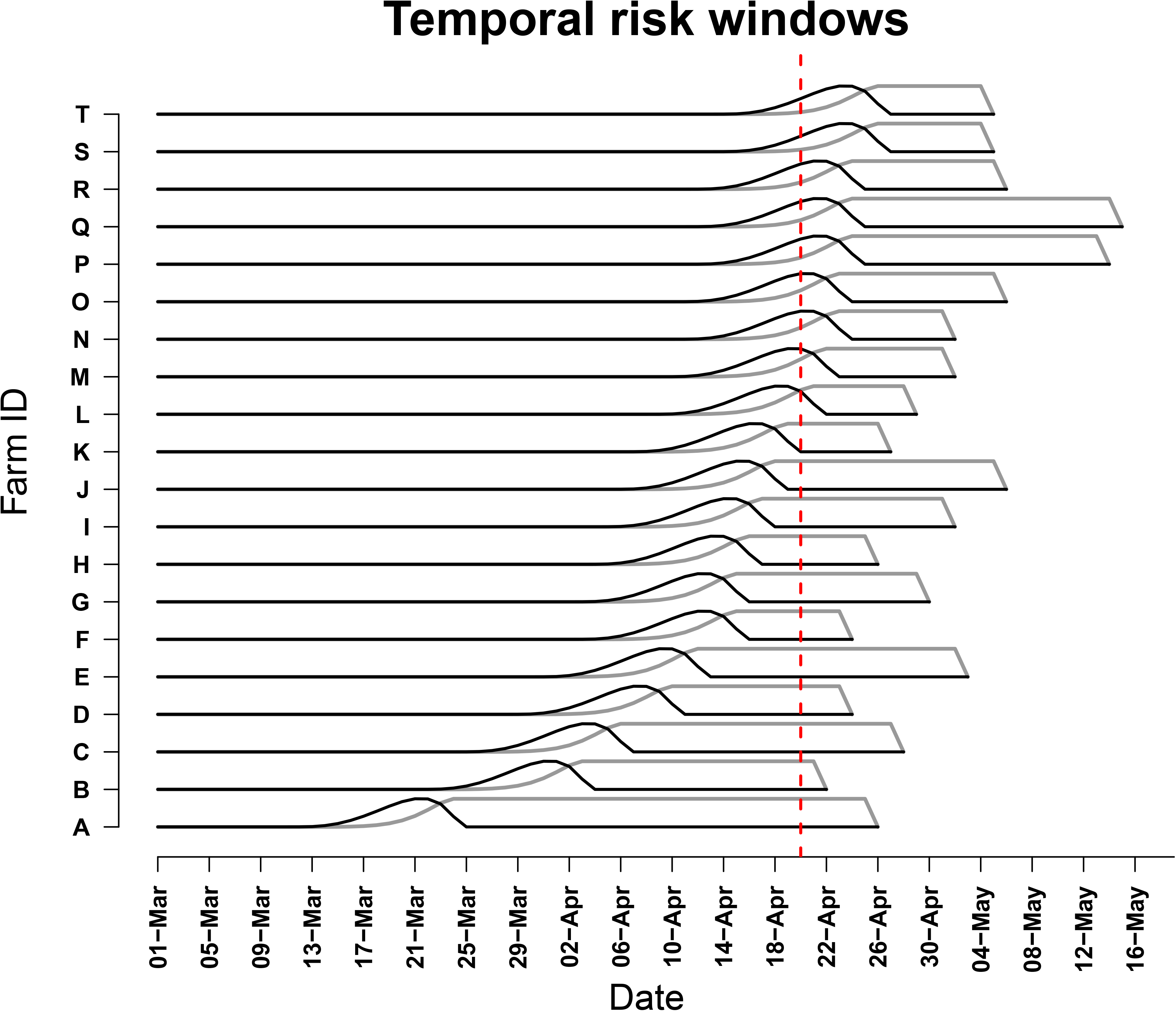
Estimated transmission windows based on Cottam’s frequentist approach for the first 20 infected premises detected for which genomic data were available in the 2010 outbreak of foot-and-mouth disease in Miyazaki Prefecture, Japan. Black lines represent most likely period of the earliest infection of an animal on each infected premises (IP), grey lines represent estimated duration of infectiousness at the premises level, tapering as culling commences. The red reference line represents the point of outbreak detection on 20 April 2010. On the most likely day that Farm B was infected, only Farm A was possibly infectious.

The posterior median estimates of the substitution rate and transition to transversion ratio inferred by SCOTTI were highly comparable to those inferred by Lau’s model, with overlapping HPDs that also encompassed the maximum likelihood value estimated using MEGA. The SCOTTI model suggested the sequence data were monophyletic (i.e., a single introduction), with only a single likely root and transmission from the original external source was estimated to have occurred 39 days prior to detection of the outbreak (i.e. on 12 March 2010). Onward transmission from the source occurred at a rate of 3.2 new infected premises per day over the course of the outbreak, with the median estimate of the number of FMD viral lineages within each farm being 19. Of those 104 IPs with genomic sequence data available, only 32 had consensus support that their proposed source was amongst those sampled and of these only 5 had >50% model support for their proposed ancestor (detailed results provided as Supplementary Materials, S6). Based on the structured coalescent transmission tree inference, there was very low likelihood that the source of infection for the first farm inferred to have been infected in this outbreak was amongst those sampled (support = 2.4%), whereas it was much more likely that the index farm’s source was amongst those sampled (support = 33.4%) and model support that the index was infected by the first farm inferred to have been infected approached consensus (42.8%).

## Discussion

Transmission network models that enable reconstruction of outbreaks hold considerable promise for informing decision-making in future outbreak responses if they are accurate, robust, reproducible, reliable and can be implemented with ease. Here, we have developed and evaluated an extended version of Lau’s systematic Bayesian inference framework incorporating additional parameters to infer farm-level effects on transmissibility and susceptibility related to the predominant species on a property and the numbers of animals kept. The modified model demonstrated improved performance across a series of varied simulated outbreaks, with overall accuracy improving by between 5 and 6%. These improvements may seem modest unless considered from the perspective that Lau’s original model was already a well-performing highly detailed inference as recently demonstrated [15] and the modified model is intended to be implemented in near-real time in outbreaks involving hundreds of infected farms, where each correctly inferred link may aid the speed of containment and subsequently greatly reduce future outbreak impacts.

The inferred transmission network for the 2010 outbreak of FMD in Japan identified all key linkages between clusters and characterised features of important farms in widespread dissemination of this outbreak. Pig farms played a vital role, with most of the farms forming hubs in the transmission network holding predominantly pigs. This has previously been identified as key to dissemination of FMD [25, 44], however, with the inclusion of additional parameters, we were able to estimate the magnitude of this effect alongside other important epidemiological and phylogenetic parameters. The five-fold increase in transmissibility of pig farms compared to farms holding predominantly cattle is biologically plausible and agrees with published accounts that, depending on FMD strain, pigs can excrete up to 100 times more airborne virus at the peak of the viraemic phase than cattle [25]. Whilst pigs may excrete more virus than ruminants, cattle on a downwind farm are more susceptible to infection via inhalation. Although pig farms tend to hold more animals, they also typically implement management measures specifically focussed on hygiene, biosecurity, ventilation, humidity and temperature control, odour and pollution reduction that would be expected to influence and often reduce the potential for disease dissemination.

The effect of numbers of animals held suggested farm size had more of an influence on farm susceptibility than transmissibility, however the HPDs of the inferred parameters representing these non-linear effects overlapped considerably. This modification was stimulated by the formulation of previous FMD models for the 2001 outbreak in the United Kingdom [24] and despite minor differences in parameterisation the estimates were all reasonably close to those fit to that prior outbreak. In some of the regions previously studied in the UK 2001 outbreak, numbers of animals held influenced transmissibility more than susceptibility, but the finding was not consistent. Such differences likely relate to differences in the predominance of sheep versus pigs in different regions and their differing influences on transmission. In their analysis, Tildesley and colleagues (2008) included species-specific parameters to represent the nonlinear influence of numbers of animals held. When we attempted to include such species-specific parameters in the modification to Lau’s approach, this led to over-parameterisation and presumed identifiability issues impacting on MCMC chain mixing and convergence. We therefore settled for a single parameter for each effect, assuming that species-specific effects should be well represented by the specific farm-level susceptibility and transmissibility terms.

The inferred farm-level incubation period in the 2010 FMD outbreak in Japan of 2–14 days corresponds very closely with previously published data [25, 45]. Interestingly, at the farm-level, the median inferred incubation period was 1 day shorter than the median latent period. This finding is consistent with an experimental study where the relationship between onset of infectiousness was based on directly demonstrating FMD transmission to another animal [46]. In contrast, many studies that have considered onset of infectiousness at the farm-level based on proxy measures (such as detection of virus in blood, nasal fluid and/or oesophageal-pharyngeal fluid) [45] may have underestimated the duration of the latent period [46]. Whilst individual animals have been shown to excrete FMD virus 1–2 days before onset of clinical signs [47–49], this depends on dose and FMD virus strain, and there is marked individual variability in the onset of early clinical signs in pigs and cattle. It is important to note that the unit of interest in the present analysis is the farm and these epidemiological parameters are therefore observed at the farm-level, whereas most studies of the timing of onset of infectiousness and clinical signs focus on the animal-level. Also, the observed epidemiological data that informed our inferences were from field observations, rather than based on experimentation, and thereby include a certain level of uncertainty. Nonetheless, these epidemiological parameters are very helpful for informing disease response activities (quarantine periods, surveillance and contact-tracing windows), and estimates from observed outbreak such as those presented here are vital for parameterising FMD simulation modelling. Similarly, the farm-level infectious period is a very important parameter, seemingly intuitive but given all the factors at play difficult to interpret. Often, as in the present analysis, the farm-level infectious period is cut short by culling and other disease control activities. In the 2010 outbreak of FMD in Japan, targeted vaccination was only implemented for 5 days at the peak of the outbreak [17], so was not considered to have had a major impact on the inference of epidemiological parameters.

With data augmenting MCMC approaches, as implemented here, reconstructing such outbreaks need not be completed years after the outbreaks are over. It is a primary intention of the design of these models that they be implemented to inform ongoing disease responses. Indeed, these models are presently being implemented in near-real time to inform the ongoing outbreak of *Mycoplasma bovis* in New Zealand [50]. As detailed in the present analysis, these models provide statistically justifiable inference of which premises were primary sources in an outbreak and the timing of exposure at those farms. This can greatly inform targeting of contact-tracing windows and farmer interviews to high-risk periods and help identify undetected sources of such outbreaks before further clusters can be seeded. An active area of further research includes incorporating contact-tracing and other animal movement data into this model. Further areas for development include refining the representation of genomic evolution through the implementation of within-host dynamics such as has been implemented in other transmission network models [10] and formally predicting undetected infections with Reversible-Jump MCMC or related methods [51].

The original attempts at FMD outbreak transmission network modelling have largely focussed on small subsets of large outbreaks [10, 12, 16, 52]. With the present modified formulation, we have demonstrated inference for outbreaks involving up to 400 premises, and with typically available parallel computing infrastructure it presently appears feasible to run inferences for outbreaks of over 500 premises with some further efficiencies in coding. The present analysis was limited in the number of simulations that could be feasibly undertaken for model verification and pseudo-validation. However, we consider the additional gain in information will be modest with further testing on substantially increased numbers of simulation runs. In the present analysis, all models had difficulties inferring secondary transmission rates when these were very low. The best-performing model was again that with the modification to incorporate farm-level effects on transmissibility and susceptibility. The low value for *β* tested in verification run J3 was perhaps unrealistic being 100 times below the inferred values based on the actual outbreak data from the 2010 outbreak in Japan. The mutation rate appears to be underestimated by all forms of the Lau model. This is not a major concern, as the primary purpose of this model is to infer the transmission network. More purposeful phylogenetic tools, such as BEAST and associated packages [35, 53], are preferable when the primary aim is estimation of such phylogenetic parameters and more sophisticated models including additional complexities such as within-host diversity are available. Nonetheless the mutation rates inferred by the modified Lau model overlapped with those of the SCOTTI model implemented in BEAST2.

The present analysis was limited in the number of simulations that could be feasibly undertaken. However, we consider the additional gain in information will be modest with further testing on substantially increased numbers of simulation runs. In the present analysis, all models had difficulties inferring secondary transmission rates when these were very low. The best-performing model was that with the modification to incorporate farm-level covariates.

There was poor agreement between the transmission networks inferred by SCOTTI and the Lau modified model. Reasons for differences in transmission network inferences include different underlying likelihood formulations and data requirements. Specifically, the Lau model infers sequences for known IPs for which genomic data is unavailable and incorporates terms that account for the spatial relationships between infected premises. For four IPs that formed an isolated cluster in Ebino, in the far West of Miyazaki Prefecture, the sources inferred by Lau’s modified model agreed very closely with epidemiological field data whereas the sources for all four of these premises inferred by SCOTTI were inferred to be over 60 km away. Whilst at least one of these premises is likely to have been infected from the main focus of infection to the East, it is highly unlikely that all four were infected in independent introductions. Considered together, the inferences of Lau and SCOTTI’s models provide a reasonably complete epidemiological and phylogenetic inference for the Japanese outbreak.

## Conclusions

Extending Lau’s systematic Bayesian inference framework to incorporate additional parameters representing predominant species and numbers of animals held on a farm resulted in improvements in overall accuracy across a series of varied simulated outbreaks. Infected farms that held predominantly pigs were estimated to have five times the transmissibility of infected cattle farms and be 49% less susceptible. The farm-level incubation period was estimated to be 1 day shorter than the latent period, suggesting a small window following onset of clinical signs to target interventions may substantially reduce the risk of onwards transmission in future outbreaks.

## Acknowledgements

This research was supported by an Australian Research Council Discovery Early Career Researcher Award (project number DE160100477) and by the Japanese Ministry of Agriculture, Forestry and Fisheries (Management Technologies for the Risk of Introduction of Livestock Infectious Diseases and Their Wildlife-borne Spread in Japan, FY2018-2022). Components of this research were undertaken on The University of Melbourne’s High Performance Computing system SPARTAN [54]. The AADIS model was initially developed by the Australian Government Department of Agriculture and Water Resources in collaboration with the University of New England and has kindly been made available to support this research. The funders had no role in study design, data collection and analysis, decision to publish or preparation of the manuscript. The authors would also like to acknowledge helpful comments on the research provided by colleagues in the National Institute of Animal Health, Japan and the Asia-Pacific Centre for Animal Health (University of Melbourne).

## Author contributions statement

All authors were involved in design of the study. SF and YH conceived the modified model formulation with input from ML and RB. SF wrote the main manuscript text and implemented all analyses. All authors reviewed and commented on the manuscript.

## Supplementary Materials

**S1: Simulated outbreak datasets: data and parameterisation.**

**S2: Comparison of the accuracy of the original model and modifications of Lau’s joint Bayesian transmission network inference for inferring sources for simulated outbreaks of foot-and-mouth disease in Japan and Australia.**

**S3: Comparison of the accuracy of the original model and modifications of Lau’s joint Bayesian transmission network inference for inferring epidemiological and phylogenetic parameters for simulated outbreaks of foot-and-mouth disease in Japan and Australia.** Model formulations abbreviated as follows: orig = original; mod = modified; mod-n = modified-normalised.

**S4: Nucleotide substitution model fit for genomic data from the 2010 outbreak of foot-and-mouth disease in Japan.**

**S5: Lau model (original and modified-normalised) inferred transmission networks and estimates for the 2010 outbreak of foot-and-mouth disease in Miyazaki Prefecture, Japan.**

