## Supplementary Materials S1-S5 for "Transmission network reconstruction for foot-and-mouth disease outbreaks incorporating farm-level covariates"

**S1 Simulated outbreak datasets: data and parameterisation**

All simulated datasets are available here: https://doi.org/10.26188/5cf5e3af414a8

In accordance with data-sharing agreements for AADIS and the Miyazaki 2010 FMD outbreak dataset, random noise has been added to the coordinates of the farms in the datasets made publicly available.

**Sellke threshold simulated FMD outbreaks in Japan (runs J1–J3)**

| **Run** | **J1** | **J2** | **J3** | **Reference/comment** |
|---|---|---|---|---|
| $n$ | 100 | 200 | 400 | Assumed outbreak size |
| *n(infected)* | 80 | 166 | 200 | Testing variety of scenarios |
| $nt$ | 7667 | 7667 | 7667 | Genome length (nucleotides) |
| $t\_max$ | 100 | 100 | 100 | Maximum length of outbreak (days) |
| $\alpha$ | 4.00E-05 | 4.00E-05 | 5.00E-06 | Adjusted to scale background transmission |
| $\beta$ | 0.085 | 0.09 | 6.00E-04 | Adjusted to scale $\beta_{ij}$ and n(infected) |
| $\mu_1$ | 2.00E-05 | 1.50E-05 | 3.00E-05 | Cottam et al (2008) |
| $\mu_2$ | 2.00E-06 | 1.00E-06 | 5.00E-06 | Cottam et al (2008) & Juleff et al (2013) |
| $a$ | 8 | 8 | 8 | Alexandersen et al (2003) & Haydon et al (2003) |
| $b$ | 0.5 | 0.5 | 0.5 | |
| $c$ | 15 | 12 | 21 | Little prior information |
| $\kappa$ *(power law)* | 1.7 | 1 | 2 | Bouma et al (2003) |
| $p$ | 0.1 | 0.1 | 0.2 | Little prior information |
| $\phi_{pigs}$ | 20 | 4 | 10 | Alexandersen et al (2003) |
| $\phi_{other}$ | 3 | 3 | 1 | |
| $\rho_{pigs}$ | 0.1 | 0.1 | 0.5 | |
| $\rho_{other}$ | 0.5 | 0.5 | 0.1 | |
| $\nu$ | 0.1 | 0.1 | 0.8 | |
| $\tau$ | 0.03 | 0.03 | 0.01 | |
| *Proportion of cattle farms* | 0.33 | 0.6 | 0.5 | |
| *Proportion of pig farms* | 0.33 | 0.1 | 0.3 | |
| *Proportion of other farms* | 0.33 | 0.4 | 0.2 | |
| *Herd size, median (range)* | 279 (8, 7862) | 288 (11, 5130) | 277 (6, 10186) | |

**AADIS simulated FMD outbreaks in Australia (runs A1–A3)**

In brief, they included n=98, 100 and 298 infected premises, respectively, and were constructed using the Australian Animal Disease Spread (AADIS) hybrid model's baseline configuration, with movement restrictions and a stamping out only policy (i.e., no vaccination), each seeded on a large pig farm in central Victoria, Australia (Bradhurst et al., 2015). Molecular sequence evolution was forwards simulated from a most recent common ancestor at the seed premises, designated with the 7667 nucleotide whole genome consensus sequence (O/JPN/2010-6/1S) sampled from the first farm presumed to be infected in the 2010 outbreak of foot-and-mouth disease in Miyazaki Prefecture of Japan (Nishi et al., 2017). Phylogenies were simulated with VirusTreeSimulator and SeqGen version 1.3.3 (Rambaut and Grass, 1997), parameterised based on empirical observations from the 2001 outbreak of FMD in the UK (Cottam et al., 2006, Cottam et al., 2008) and 2010 outbreak in Japan (Nishi et al., 2017).

**Supplementary Materials, S2**

**Table S2: Comparison of the accuracy of inferences of Lau's joint Bayesian inference of the transmission network and two modified models, for six simulated outbreaks of foot-and-mouth disease in Japan and Australia, detailed by run.**

| **Run**[a] | **Model**[b] | **Accuracy**[c] Overall (%) | >50% support (%) | >80% support (%) |
|---|---|---|---|---|
| J1 | original | 80/100 (80) | 78/88 (89) | 62/66 (94) |
| | extended | 85/100 (85) | 83/92 (90) | 76/79 (96) |
| | extended (norm) | 84/100 (84) | 82/91 (90) | 77/81 (95) |
| J2 | original | 159/200 (80) | 149/167 (89) | 126/129 (98) |
| | extended | 170/200 (85) | 158/174 (91) | 134/138 (97) |
| | extended (norm) | 167/200 (84) | 156/173 (90) | 135/142 (95) |
| J3 | original | 357/400 (89) | 334/348 (96) | 287/289 (99) |
| | extended | 376/400 (94) | 357/367 (97) | 326/327 (99) |
| | extended (norm) | 372/400 (93) | 359/370 (97) | 331/332 (99) |
| A1 | original | 77/98 (79) | 71/83 (86) | 54/60 (90) |
| | extended | 83/98 (85) | 81/88 (92) | 63/69 (91) |
| | extended (norm) | 84/98 (86) | 80/88 (91) | 56/59 (95) |
| A2 | original | 74/100 (74) | 70/93 (75) | 48/57 (84) |
| | extended | 77/100 (77) | 74/92 (80) | 53/56 (95) |
| | extended (norm) | 75/100 (75) | 74/98 (76) | 57/67 (85) |
| A3 | original | 222/298 (75) | 206/248 (83) | 152/169 (90) |
| | extended | 227/298 (76) | 217/252 (86) | 164/173 (95) |
| | extended (norm) | 226/298 (76) | 215/254 (85) | 162/171 (95) |

norm = normalised; IP = infected premises. [a] Runs J1, J2 and J3 were FMD outbreaks in Miyazaki Prefecture of Japan simulated in the same framework as the Extended model. Runs A1, A2, A3 were FMD outbreaks in south-eastern Australia simulated in using the Australian Animal Disease Simulation (AADIS) model (Bradhurst et al., 2015). [b] The original model is as described in (Lau et al., 2015, Firestone et al., provisionally accepted for publication). The extended model incorporated additional terms for farm level transmissibility and susceptibility based on farm type and number of animals. The extended (norm) model incorporated the same terms, with normalisation by mean transmissibility and susceptibility of all farms. [c] Accuracy was defined as the proportion of IPs for which the model-predicted most likely source (highest likelihood or most posterior support) was the true source. The denominator for accuracy at >50% and >80% support includes only those IPs for which the model-predicted most likely source attained that level of likelihood or posterior support.

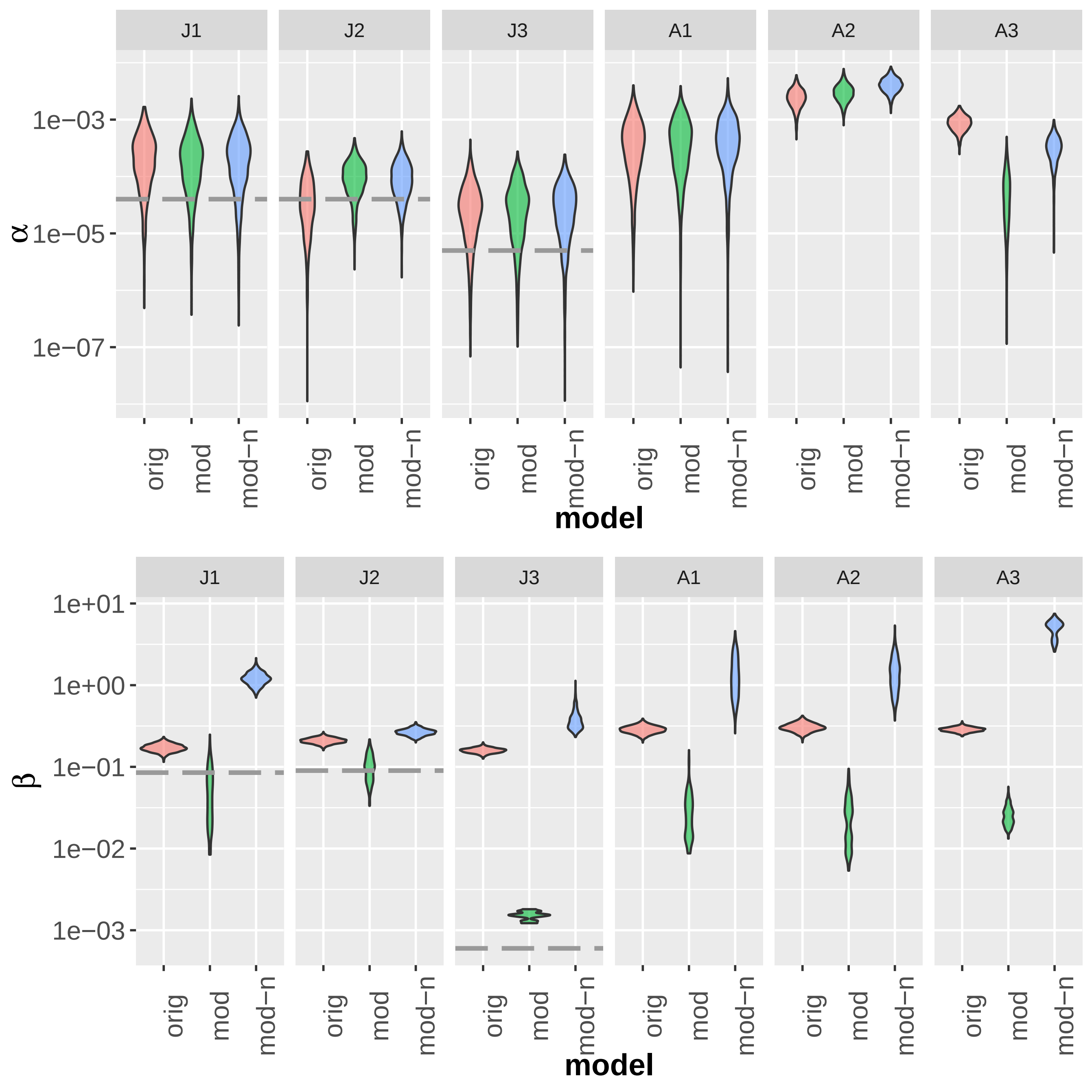
J1
J2
J3
A1
A2
A3
1e−03
1e−05
1e−07
α
orig
mod
mod−n
model
J1
J2
J3
A1
A2
A3
1e+01
1e+00
1e−01
1e−02
1e−03
β
orig
mod
mod−n
model

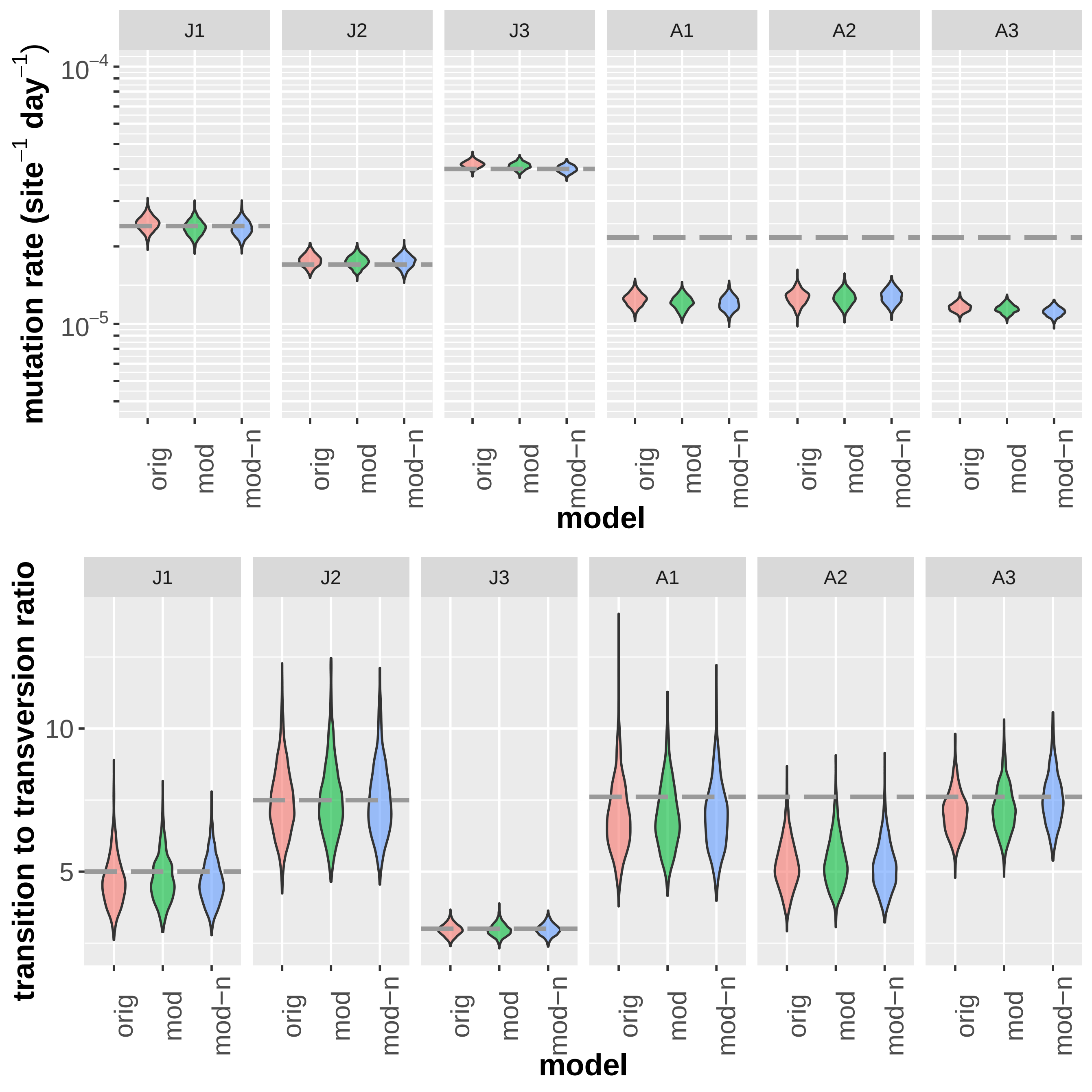

J1
J2
J3
A1
A2
A3
mutation rate (site$^{-1}$ day$^{-1}$)
$10^{-4}$
$10^{-5}$
orig
mod
mod−n
model
transition to transversion ratio
10
5
orig
mod
mod−n
model

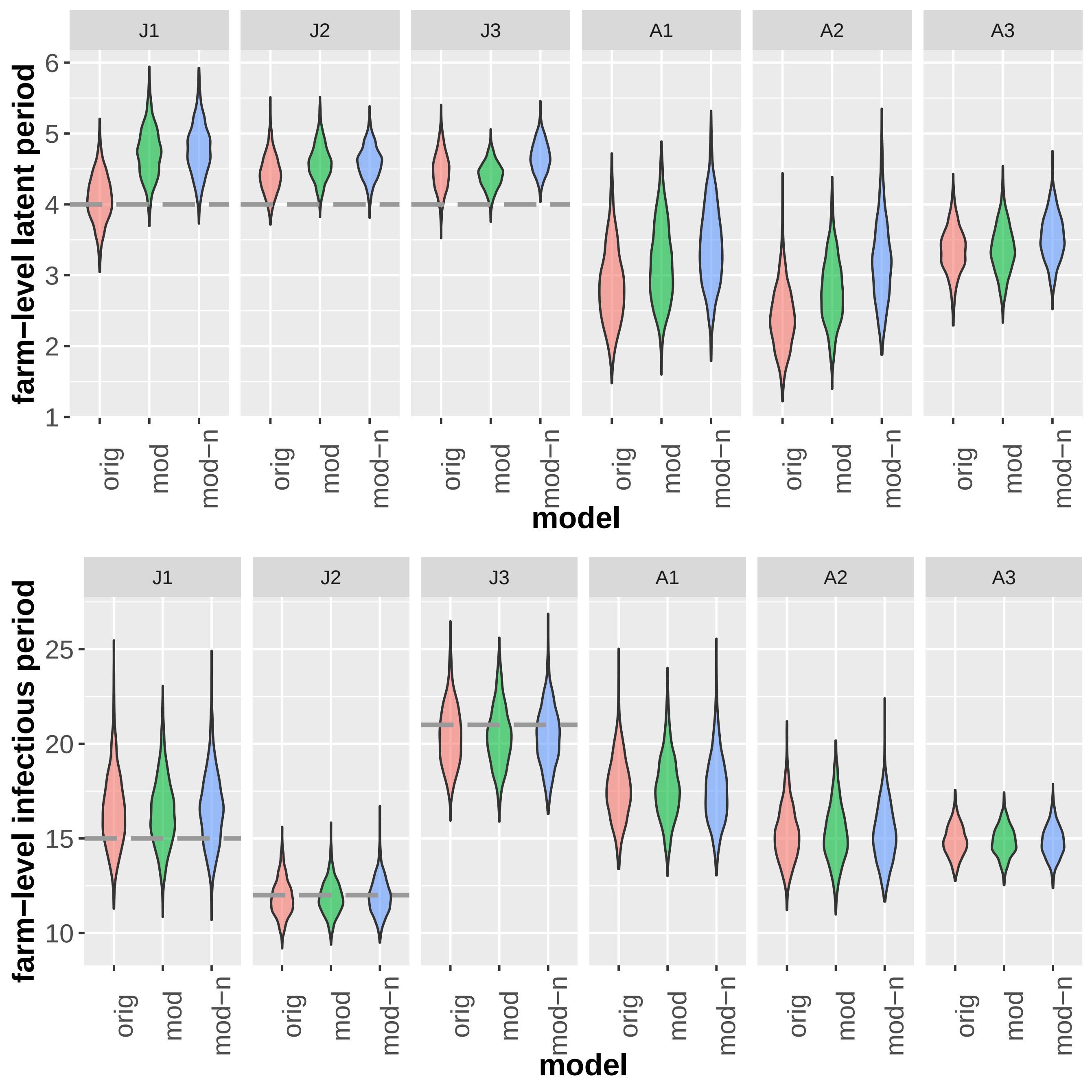
J1
J2
J3
A1
A2
A3
farm−level latent period
1
2
3
4
5
6
orig
mod
mod−n
model
farm−level infectious period
10
15
20
25
model

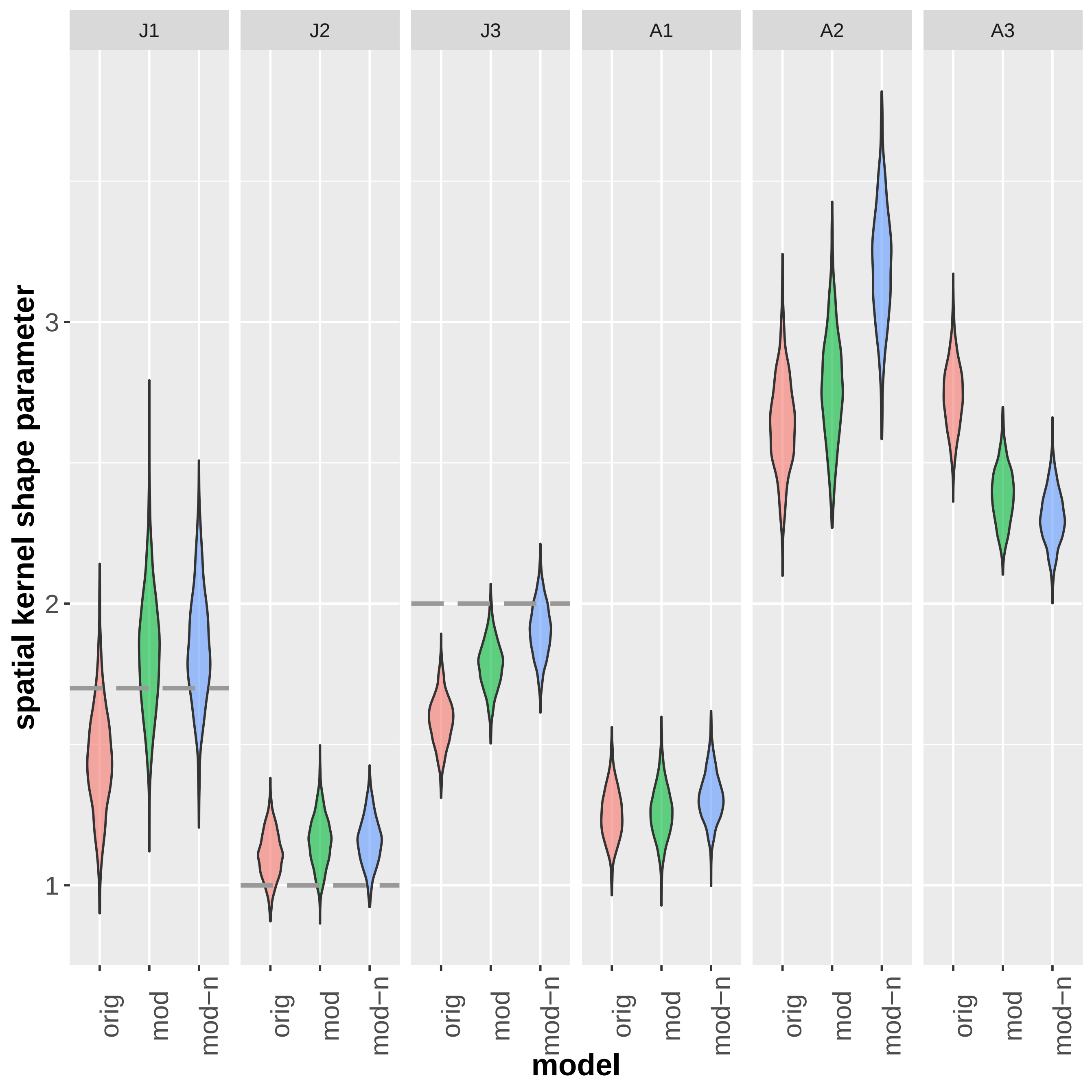

J1
J2
J3
A1
A2
A3
spatial kernel shape parameter
3
2
1
orig
mod
mod−n
model

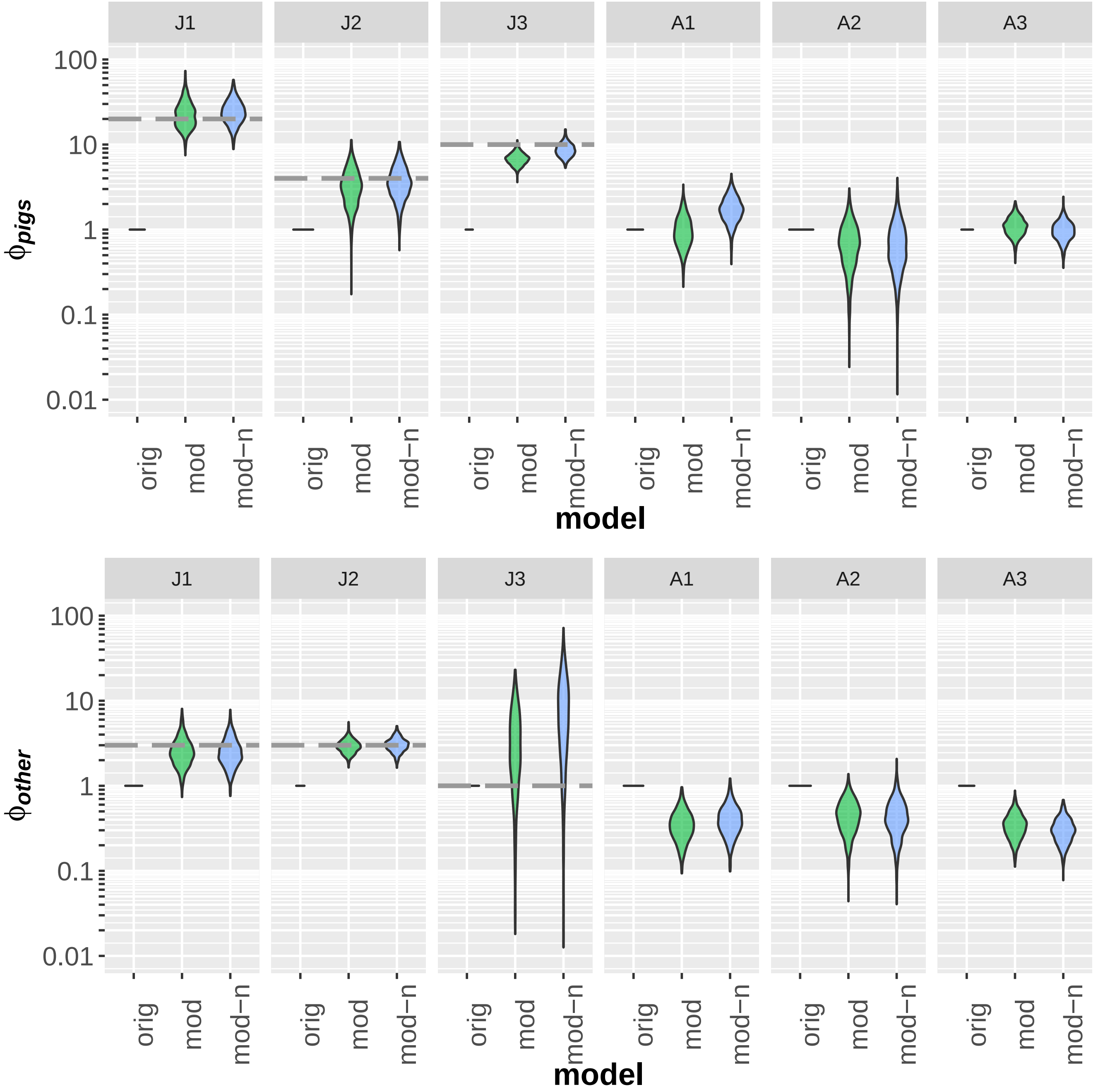
J1
J2
J3
A1
A2
A3
100
10
1
0.1
0.01
$\phi_{pigs}$
orig
mod
mod−n
model
$\phi_{other}$

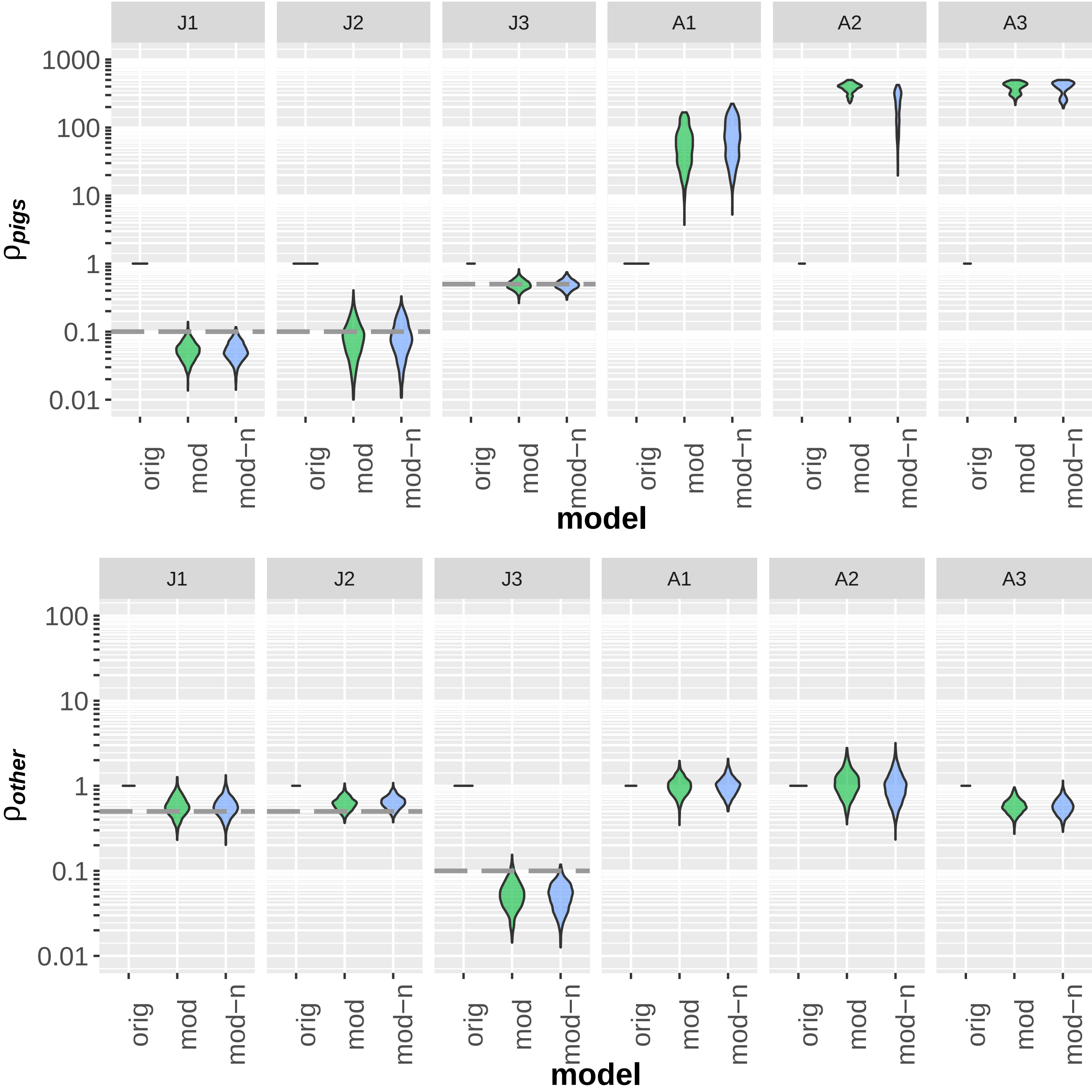

J1
J2
J3
A1
A2
A3
1000
100
10
1
0.1
0.01
ρpigs
orig
mod
mod−n
model
100
10
1
0.1
0.01
ρother

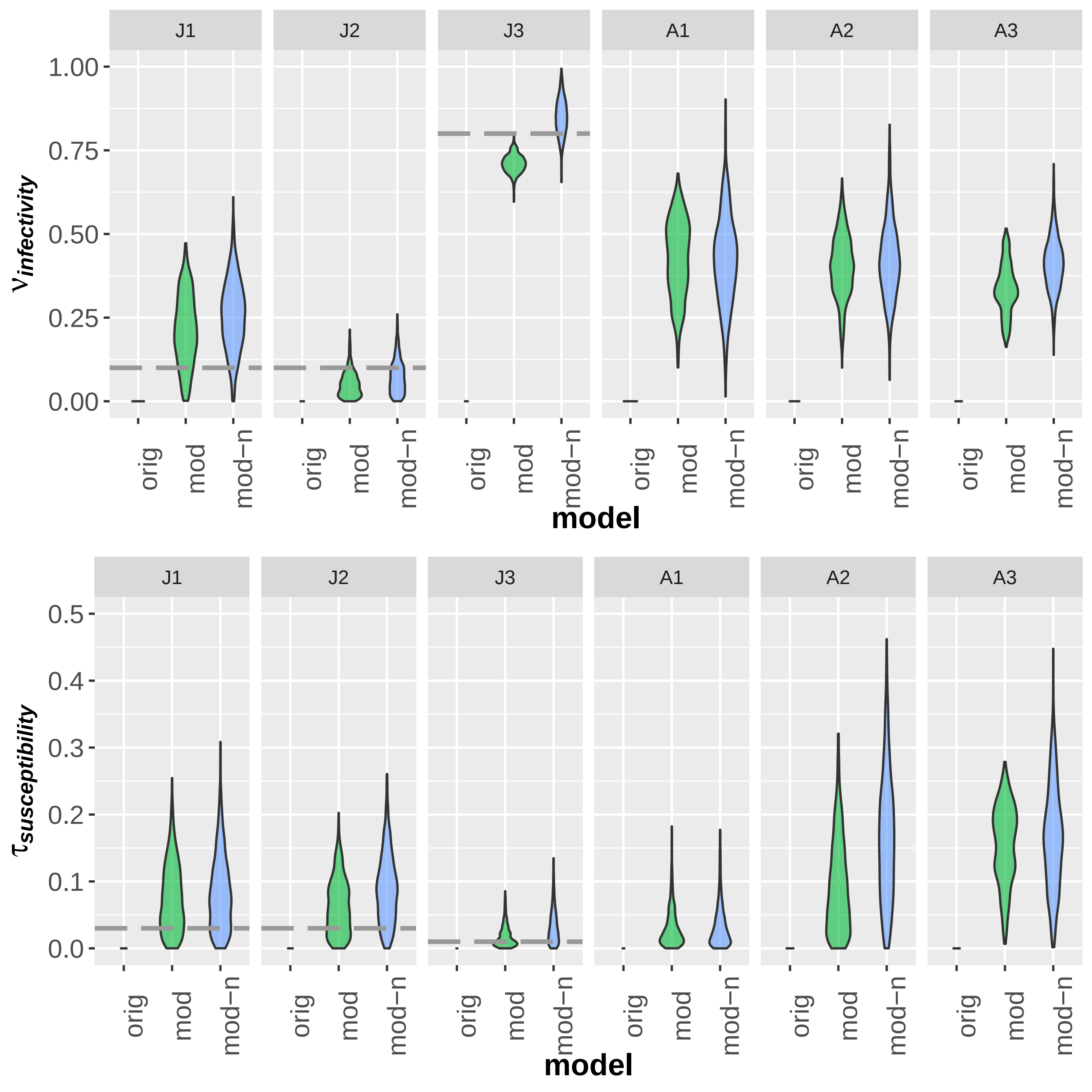
J1
J2
J3
A1
A2
A3
1.00
0.75
0.50
0.25
0.00
$\nu_{infectivity}$
orig
mod
mod−n
model
0.5
0.4
0.3
0.2
0.1
0.0
$\tau_{susceptibility}$
model

**S4 Nucleotide substitution model fit, FMD JPN 2010**

| Model | Parameters | BIC | AICc | lnL | I | G | TsTv | Freq A | Freq T | Freq C | Freq G |
|---|---|---|---|---|---|---|---|---|---|---|---|
| TN93+G | 211 | 28855.92837 | 26413.74145 | -12995.81381 | n/a | 0.130200216 | 9.076621971 | 0.25267511 | 0.207517529 | 0.282105132 | 0.25770223 |
| TN93+G+I | 212 | 28864.53158 | 26410.77085 | -12993.32797 | 0.658211802 | 1.128512764 | 9.078792105 | 0.25267511 | 0.207517529 | 0.282105132 | 0.25770223 |
| TN93+I | 211 | 28874.867 | 26432.68008 | -13005.28312 | 0.48313269 | n/a | 9.054655315 | 0.25267511 | 0.207517529 | 0.282105132 | 0.25770223 |
| HKY+G | 210 | 28884.22187 | 26453.60876 | -13016.748 | n/a | 0.091456744 | 9.078205262 | 0.25267511 | 0.207517529 | 0.282105132 | 0.25770223 |
| TN93 | 210 | 28886.71381 | 26456.1007 | -13017.99397 | n/a | n/a | 9.039801569 | 0.25267511 | 0.207517529 | 0.282105132 | 0.25770223 |
| HKY+G+I | 211 | 28891.65401 | 26449.46709 | -13013.67663 | 0.6945073 | 1.117198154 | 9.088293207 | 0.25267511 | 0.207517529 | 0.282105132 | 0.25770223 |
| T92+G | 208 | 28891.89381 | 26484.42833 | -13034.15885 | n/a | 0.098889927 | 9.05254026 | 0.230096319 | 0.230096319 | 0.269903681 | 0.269903681 |
| GTR+G | 214 | 28895.57441 | 26418.66609 | -12995.2745 | n/a | 0.130759886 | 7.514631099 | 0.25267511 | 0.207517529 | 0.282105132 | 0.25770223 |
| T92+G+I | 209 | 28899.50321 | 26480.46392 | -13031.17612 | 0.688643473 | 1.117986426 | 9.056165217 | 0.230096319 | 0.230096319 | 0.269903681 | 0.269903681 |
| GTR+G+I | 215 | 28904.0168 | 26415.53469 | -12992.70825 | 0.65793394 | 1.129782672 | 7.538650152 | 0.25267511 | 0.207517529 | 0.282105132 | 0.25770223 |
| HKY+I | 210 | 28906.01982 | 26475.40671 | -13027.64698 | 0.48313269 | n/a | 9.049551648 | 0.25267511 | 0.207517529 | 0.282105132 | 0.25770223 |
| GTR+I | 214 | 28911.50824 | 26434.59991 | -13003.24141 | 0.48313269 | n/a | 8.178632638 | 0.25267511 | 0.207517529 | 0.282105132 | 0.25770223 |
| T92+I | 208 | 28913.13531 | 26505.66983 | -13044.7796 | 0.48313269 | n/a | 9.042822042 | 0.230096319 | 0.230096319 | 0.269903681 | 0.269903681 |
| HKY | 209 | 28918.7708 | 26499.73151 | -13040.80991 | n/a | n/a | 9.03912873 | 0.25267511 | 0.207517529 | 0.282105132 | 0.25770223 |
| T92 | 207 | 28925.68408 | 26529.79243 | -13057.84143 | n/a | n/a | 9.03906144 | 0.230096319 | 0.230096319 | 0.269903681 | 0.269903681 |
| GTR | 213 | 28926.36894 | 26461.03442 | -13017.45921 | n/a | n/a | 7.460939873 | 0.25267511 | 0.207517529 | 0.282105132 | 0.25770223 |
| K2+G | 207 | 28926.5078 | 26530.61616 | -13058.25329 | n/a | 0.096171882 | 9.054583311 | 0.25 | 0.25 | 0.25 | 0.25 |
| K2+G+I | 208 | 28934.03521 | 26526.56974 | -13055.22956 | 0.690861506 | 1.115134899 | 9.059015367 | 0.25 | 0.25 | 0.25 | 0.25 |
| K2+I | 207 | 28947.95835 | 26552.06671 | -13068.97857 | 0.48313269 | n/a | 9.043104122 | 0.25 | 0.25 | 0.25 | 0.25 |
| K2 | 206 | 28960.5753 | 26576.25749 | -13082.07449 | n/a | n/a | 9.039041382 | 0.25 | 0.25 | 0.25 | 0.25 |
| JC+G | 206 | 29309.27585 | 26924.95803 | -13256.42476 | n/a | 0.096748703 | 0.5 | 0.25 | 0.25 | 0.25 | 0.25 |
| JC+G+I | 207 | 29316.84345 | 26920.9518 | -13253.42112 | 0.690282044 | 1.116562183 | 0.5 | 0.25 | 0.25 | 0.25 | 0.25 |
| JC+I | 206 | 29330.67429 | 26946.35648 | -13267.12398 | 0.48313269 | n/a | 0.5 | 0.25 | 0.25 | 0.25 | 0.25 |
| JC | 205 | 29343.27176 | 26970.52778 | -13280.21016 | n/a | n/a | 0.5 | 0.25 | 0.25 | 0.25 | 0.25 |

All positions containing gaps and missing data were eliminated. There were a total of 7559 positions in the final dataset. Evolutionary analyses were conducted in MEGA7[1,2].
Abbreviations: Bayesian Information Criterion (BIC); Akaike Information Criterion corrected (AICc); Maximum negative log Likelihood value (lnL); Proportion invariant (I); Discretised Gamma distribution shape parameter (G); Transition to transversion ratio (TsTv); Empirically estimated frequencies

**Lau model (original) inferred transmission network in arbitrary space, differences from modified network in red.**

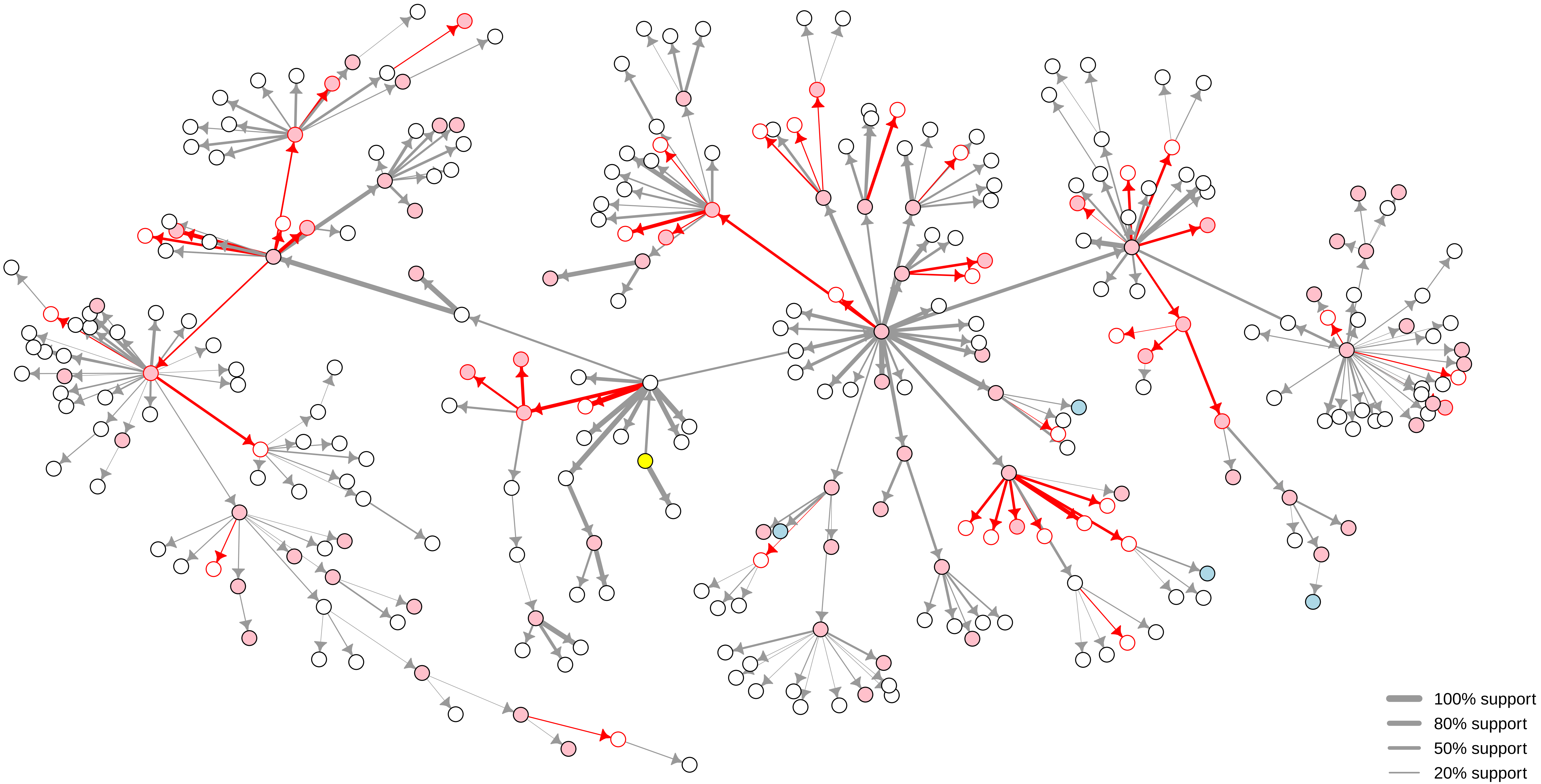

**Lau model (modified, normalised) inferred transmission network in arbitrary space, differences from modified network in red.**

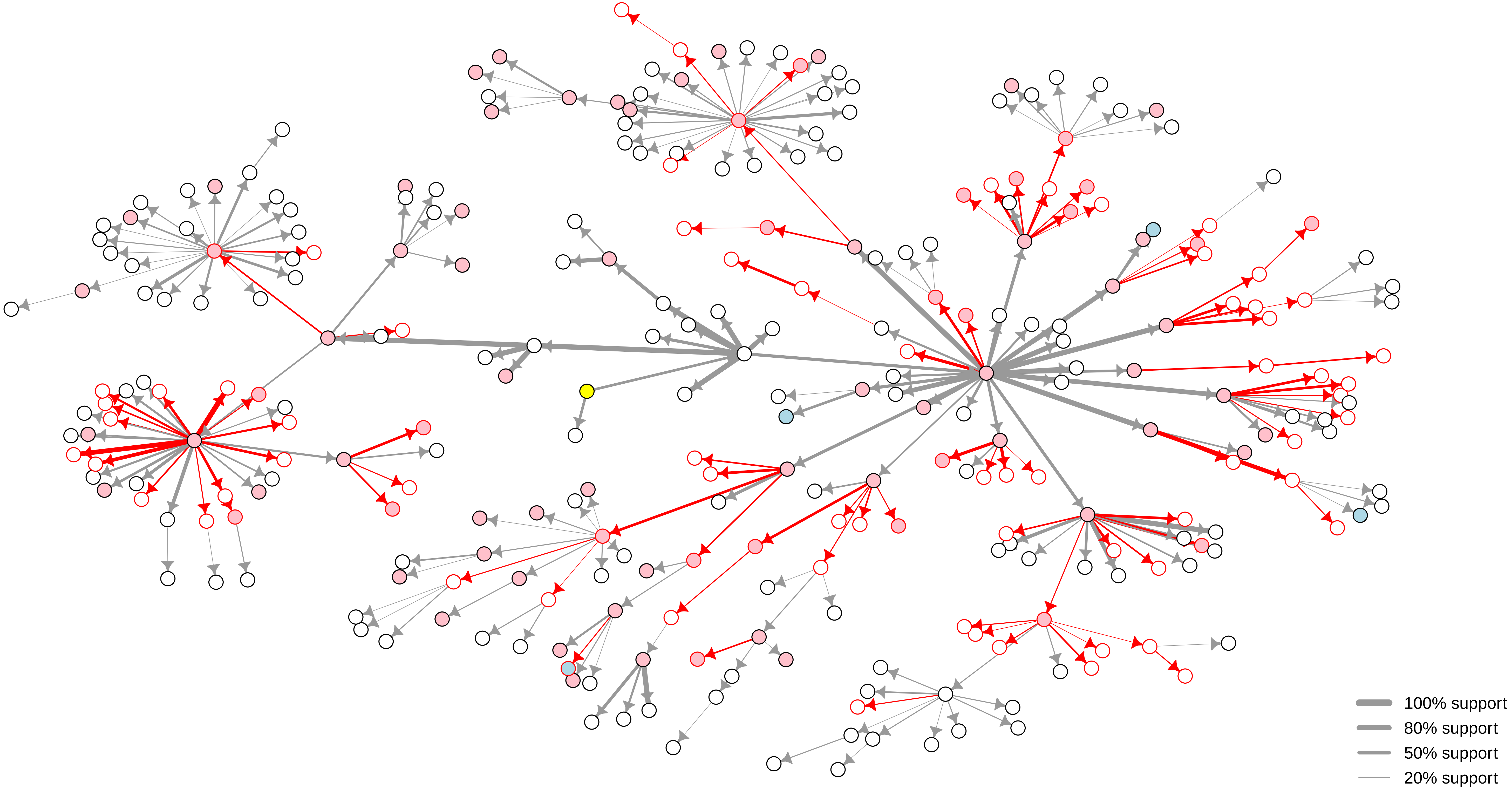
